# Induction of Ferroptosis by an Amalgam of Extracellular Vesicles and Iron Oxide Nanoparticles Overcomes Cisplatin Resistance in Lung Cancer

**DOI:** 10.1101/2024.08.19.608664

**Authors:** Anjugam Paramanantham, Rahmat Asfiya, Yariswamy Manjunath, Lei Xu, Grace McCully, Siddharth Das, Hu Yang, Jussuf T. Kaifi, Akhil Srivastava

**Affiliations:** Department of Pathology and Anatomical Sciences, University of Missouri School of Medicine, Columbia, Missouri 65212, USA; Surgery, University of Missouri School of Medicine, Columbia, Missouri 65212, USA; Ellis Fischel Cancer Center, University of Missouri School of Medicine, Columbia, Missouri 65212, USA; Harry S. Truman Memorial Veterans’ Hospital, Columbia, Missouri 65201, USA; Linda and Bipin Doshi Department of Chemical and Biochemical Engineering, Missouri University of Science and Technology, Rolla, Missouri 65409, USA

**Keywords:** Extracellular vesicles, drug delivery, ferroptosis, iron oxide nanoparticles, cisplatin resistance

## Abstract

Extracellular vesicles (EVs) hold potential as effective carriers for drug delivery, providing a promising approach to resolving challenges in lung cancer treatment. Traditional treatments, such as with the chemotherapy drug cisplatin, encounter resistance in standard cell death pathways like apoptosis, prompting the need to explore alternative approaches. This study investigates the potential of iron oxide nanoparticles (IONP) and EVs to induce ferroptosis—a regulated cell death mechanism—in lung cancer cells. We formulated a novel EV and IONP-based system, namely ‘ExoFeR’, and observed that ExoFeR demonstrated efficient ferroptosis induction, evidenced by downregulation of ferroptosis markers (xCT/SLC7A11 and GPX4), increased intracellular and mitochondrial ferrous iron levels, and morphological changes in mitochondria. To enhance efficacy, tumor-targeting transferrin (TF)-conjugated ExoFeR (ExoFeR^TF^) was developed. ExoFeR^TF^ outperformed ExoFeR, exhibiting higher uptake and cell death in lung cancer cells. Mechanistically, nuclear factor erythroid 2-related factor 2 (Nrf2)—a key regulator of genes involved in glutathione biosynthesis, antioxidant responses, lipid metabolism, and iron metabolism—was found downregulated in the ferroptotic cells. Inhibition of Nrf2 intracellular translocation in ExoFeR^TF^-treated cells was also observed, emphasizing the role of Nrf2 in modulating ferroptosis-dependent cell death. Furthermore, ExoFeR and ExoFeR^TF^ demonstrated the ability to sensitize chemo-resistant cancer cells, including cisplatin-resistant lung cancer patient-derived tumoroid organoids. In summary, ExoFeR^TF^ presents a promising and multifaceted therapeutic approach for combating lung cancer by intrinsically inducing ferroptosis and sensitizing chemo-resistant cells.

## 1. Introduction

Extracellular vesicles (EVs) are natural nanocarriers in the body, and they enable cell-to-cell communication by transporting macromolecules. Encapsulating various biological materials like nucleic acids, lipids, RNAs, and proteins within a lipid bilayer (Kanada et al., 2015; Zomer et al., 2015), EVs exhibit biocompatibility, low immunogenicity, and inherent targeting abilities, rendering them a promising delivery system for a wide range of diseases (Alvarez-Erviti et al., 2011; Chinnappan et al., 2020; Mentkowski & Lang, 2019; Srivastava et al., 2016; Tian et al., 2014).

Lung cancer remains a formidable global health challenge, claiming approximately 2.2 million new cases and 1.76 million deaths per year (Siegel et al., 2023). Traditional cancer treatments like immunotherapy, chemotherapy, and radiation aim to eliminate cancer cells by restoring the dysregulated apoptotic mechanism (Carneiro & El-Deiry, 2020). Cisplatin (cis-diamminedichloroplatinum II; CDDP), a traditional chemotherapy agent, has been the first-line treatment for decades in lung cancer. It primarily induces apoptosis; however, over time, cancer cells develop resistance by preventing the formation of platinum-DNA adducts and acquiring mutations that resist apoptosis, leading to the failure of therapy (Carneiro & El-Deiry, 2020; Galluzzi et al., 2012).

In such circumstances, an alternate form of regulated cell death (RCD) could be crucial for overcoming therapeutic resistance. Ferroptosis is a recently discovered iron ion-dependent, non-apoptotic form of RCD, which causes peroxidation of lipids in the cell membrane, causing membrane disintegration, and eventually leading to cell death (Dixon et al., 2012). Compounds like Erastin (Dixon et al., 2012), Sorafenib (Dixon et al., 2014), RSL3 (Yang & Stockwell, 2008), and Sulfasalazine (H. Yu et al., 2019) have been explored as ferroptosis inducers (FINs). Despite demonstrating efficacy in triggering ferroptosis, these molecules faced challenges in *in vivo* studies due to low stability and toxicity (Yang et al., 2014; Zou & Schreiber, 2020). Iron nanoparticles (IONP) are potent FIN and thus offer a promising therapeutic strategy for treating resistant tumors (S. E. Kim et al., 2016; Z. Li et al., 2022). However, the application of bare IONPs as a therapeutic modality in the form of FIN remains challenging due to disadvantages such as poor biodistribution, non-targeted cytotoxicity, and immunogenic incompatibilities.

Thus, taking advantage of EVs’ properties as natural cargo carriers, we engineered EVs to incorporate IONP and use the system to cause intrinsic ferroptosis induction in tumor cells. We coined the term ExoFeR for this novel EV-IONP complex and tested its capability as a robust approach to combat treatment resistance in lung cancer.

Transferrin (TF), a serum protein, is commonly utilized as a cancer-targeting agent due to its ability to bind to transferrin receptors overexpressed on the cell surface of proliferating cancer cells (Citores et al., 2002; Gatter et al., 1983). This makes TF an attractive choice for a targeted drug delivery system in cancer therapy, meeting both safety and selectivity criteria. TF was added to the ExoFeR to create an ExoFeR^TF^ complex that can induce ferroptosis specifically in cancer cells. This system is effective in inducing ferroptosis, as shown by the downregulation of ferroptosis markers, increase in iron levels, and changes in mitochondria structure. The ExoFeR^TF^ has higher uptake and can induce higher levels of tumor cell death when compared to ExoFeR. The study also identified that the downregulation of Nrf2 in ferroptotic cells and the inhibition of Nrf2 intracellular translocation in ExoFeR^TF^-treated cells play a significant role in modulating ferroptosis-dependent cell death. Furthermore, ExoFeR and ExoFeR^TF^ have the potential to sensitize chemo-resistant cancer cells, including cisplatin-resistant lung cancer patient-derived tumor organoids (PDTOs), making it a novel promising multifaceted therapeutic approach for lung cancer treatment.

## 2. Materials and methods

### 2.1. Cell culture

The NCI-A549, NCI-HCC827 human lung cancer cell lines, and normal lung fibroblast cells NCI-MRC-9 were acquired from the American Type Culture Collection (ATCC). NCI-MRC-9 was cultured in Minimum Essential Medium (MEM), while NCI-A549 was cultured in Dulbecco’s Modified Eagle Medium (DMEM) (Hyclone, Marlborough, MA, USA) and NCI-HCC827 cells were cultured in Roswell Park Memorial Institute (RPMI) medium (Hyclone, Marlborough, MA, USA). The A2780 human ovarian carcinoma cell lines and their cisplatin-resistant subclone, A2780-CIS, were acquired from Sigma Aldrich and were cultured in Roswell Park Memorial Institute (RPMI) medium (Hyclone, Marlborough, MA, USA). A concentration of 1 μM cisplatin was added to the culture medium to maintain the cells. The culture medium contained 10 % heat-inactivated (v/v) exosome-free fetal bovine serum (FBS) (Systems Biosciences, Cat. No. EXO-FBSHI-250A-1), 1 mM l-glutamine, 100 U/mL penicillin, and 100 μg/mL streptomycin (Thermo Fisher Scientific, Cat. No. P4333). The cells were maintained at 37 °C in a humidified atmosphere of 95% air and 5% CO2. Routine PCR assessment with specific oligonucleotides was conducted to monitor the cells for mycoplasma contamination. GW4869 (Selleck, Cat. No. S7609) was utilized to inhibit the production of EVs.

### 2.2. Patient-derived tumor organoid (PDTO) culture

Tumor specimens obtained from non-small cell lung cancer (NSCLC) patients (MU387, MU377, MU369, and MU383) were sourced from the University of Missouri Hospital, following the approved IRB no. 2010166 and obtaining written informed consent. Upon receipt, the samples underwent a thorough washing with 1X PBS three times. Subsequently, the specimens were dissected into small pieces (1-2 mm³) using sterile instruments, then combined with 20 mL of digestion medium (DMEM/F12, 0.4% fungizone, 1% antibiotics, 500 µg/mL collagenase I, 25 µg/mL DNase I, 25 µg/mL elastase, 100 µg/mL hyaluronidase) for one hour at 37°C, 5% CO2 in an incubator with agitation (200 rpm).

Following the incubation, the suspension was filtered through 70-μm cell strainers (Corning, Cat. No. 352350), and the strained cells were centrifuged at 240 × g for 4 minutes. The resulting pellet was then resuspended in DMEM/F12-Dulbecco’s Modified Eagle Medium and Nutrient Mixture F-12 (Thermo Fisher Scientific, Cat. No. 12634010), containing bFGF 20 ng/mL (Invitrogen, Cat. No. RP-8627), EGF 50 ng/mL (Invitrogen Cat. No. RP-10927), B-27 Supplement (Invitrogen, Cat. No. 12587010), 1 × ROCK Inhibitor (Y-27632), and Penicillin-Streptomycin-Amphotericin B Suspension, followed by centrifugation at 240 × g for 4 minutes. The patient-derived cells were then suspended in ECM Gel (Sigma, Cat. No. E1270-5ML) at a concentration of 10^7^ cells/mL. The ECM gel was liquefied at 4 °C before use, and the suspension was seeded in a 24-well culture plate and incubated at 37 °C for gelation. Subsequently, 500 μL of DMEM/F12 was added to each well, and the media was changed every 4 days. Once tumoroids formed, they underwent passage through the following steps: the tumoroids with ECM gels were collected using a 1 ml tip, placed on ice for a few minutes, mixed with media, and then subjected to centrifugation at 800 × g for 4 minutes. This step was repeated to eliminate the ECM gel. Prechilled phosphate buffer saline (PBS) was added and washed to remove any residual gel. The isolated tumoroids were thoroughly washed and centrifuged at 800 × g for 5 minutes at 4 °C. These steps were iterated until all residual gel was completely removed. Purified tumoroids were then reseeded in the ECM gel in a new culture plate.

### 2.3. EVs isolation

For the isolation of EVs from MRC-9 cell lines, cell culture media was employed. MRC-9 cells were initially seeded at a density of 1×10^6^ cells/dish in T75 flasks (Cellstar Cat. No. F010013) and cultured in a conditioned medium composed of MEM with exosome-free-FBS. After a 72-hour incubation period, the medium was removed from the cells. Prior to EV isolation, the media underwent clarification through centrifugation at 2500 × g for 30 minutes (Sorvall ST4 Plus Centrifuge Series) and filtration using 0.22 µm syringe filters (Thermo Fisher Scientific, MA, USA, Cat. No. SLGVV255F). This step aimed to eliminate detached cells, large debris, and contaminants such as large particles (ectosomes, microvesicles). The supernatant obtained was then subjected to ultracentrifugation (UC) at 100,000 × g at 4 °C for 120 minutes, using a 70.2 Ti rotor (Beckman Coulter Optima XPN-80), to pellet the EVs. The resulting pellets were carefully separated from the supernatant, resuspended in 1X Phosphate Buffered Saline (PBS) (Corning, Cat. No. 21-040-CV), and subsequently stored at -80 °C.

### 2.4. EV characterization

#### 2.4.1. Nanoparticle tracking analysis system (NTA)

The NanoSight NS300 system from NanoSight in Malvern, Worcestershire, UK, was utilized to characterize the size distribution and quantity of EVs isolated through ultracentrifugation (UC). An automated syringe pump system (Harvard apparatus, Cat. No. 98-4730) with a constant flow rate of 100 was employed to inject 1 ml of purified small EVs into the system. The samples were diluted to a final volume of 1 ml in PBS. Preliminary testing was conducted to determine the optimal measurement concentrations, considering the ideal particle per frame value (20–100 particles/frame). Subsequent adjustments were made following the guidelines provided in the software handbook (NanoSight NS300 User Manual, MAN0541-01-EN-00, 2017): the camera level was increased until all particles were clearly visible without exceeding a particle signal saturation of over 20%. The moving samples in the flow cell were illuminated using a 530 nm laser, and the scattering track produced by each particle was recorded in a 60-second video using an sCMOS camera. Three similar runs were conducted to ensure statistically significant data. The acquired data was analyzed using NTA 3.2 software to obtain the final estimates of EV particle size and concentration. The NTA capture and analysis parameters were consistently maintained for all runs.

#### 2.4.2. Transmission electron microscopy (TEM)

EVs were mounted onto 300 hex mesh copper grids coated with formvar and subjected to glow discharge using the single drop method. A syringe, fitted with a new 0.2 mm filter, contained staining solutions and deionized water. Subsequently, 10 ml of the sample was applied to the grid, allowing it to settle for 5 minutes. Any remaining unsettled sample was removed by wicking with filter paper, followed by a 10-second wash with deionized water. Excess water was eliminated using filter paper, and then 10 ml of 2% Uranyl acetate was applied to the grid for 20 seconds. For IONPs and ExoFeR samples, the same procedure was followed without staining. The grid was left to dry in a desiccator and then stored in a grid storage box, ready for viewing. The grids were examined using a JEOL JEM-1400 Transmission Electron Microscope at 80 kV, equipped with a 2k AMT digital camera (JEOL, Tokyo, Japan).

### 2.5. ExoFeR complex preparation

200 µg protein equivalent of EVs isolated from MRC-9 cells were incubated with 20, 40 µg of iron oxide nanoparticles (IONPs; 10 nm size) made up to the volume of 200 µl of PBS (pH 7.4). For ExoFeR^TF^ formulation human TF (Tf, 76-81kDa MW) was first converted into thiolate Tf by treating with the iminothiolane reagent. Briefly, 6.5 mg of Tf was dissolved in 2 ml of 0.1M Na2HPO4 with 0.1M EDTA at pH 8.0 and 0.25 mg of iminothiolane added and stirred for 3 hours at 4^°^C to form the Tf-iminothiolane (Tf-SH) solution. Any unbound iminothiolane was removed using PD10 desalting column with 0.15M NaCl solution used as column equilibrating buffer and 0.1M Na2HPO4 with 0.1M EDTA at pH 7.1 as eluent buffer. Then Tf concentration was estimated by BCA protein assay and 2.7 mg of purified Tf-SH was mixed with one mg of maleimide-PEG-PE lipid (2941 MW) and allowed to react overnight at 4°C. The resulting Tf-DSPE lipid was purified by centrifugation at 16,000 x g for 15 minutes and the Tf concentration in the Tf-DSPE lipid was estimated using the BCA protein assay kit. For inserting the Tf-DSPE lipid into EVs, 200 µg protein equivalent of EVs isolated from MRC-9 cells were dispersed in PBS (pH 7.4). EVs were loaded with 4 µg of TF equivalent from a Tf-DSPE lipid complex, then placed in an incubator at 37°C for 1 hour with gentle stirring.

#### 2.5.1. Inductively coupled plasma mass spectrometry (ICP-MS)

Acid digestion was conducted for sample pretreatment. Concentrated aqua regia was used to digest the sample completely. All plasticwares used in this study were pre-cleaned by soaking in 3% HNO3 overnight prior to use. Calibration standard solutions were prepared by diluting 1,000 mg/l stock solution containing Fe ion to different concentrations using1% HNO3. Bismuth (Bi) solution (5 µg/l) was selected as the internal standard and added online during the ICP-MS analysis (NexION 2000, Perkin Elmer, USA). The method detection limits were also checked periodically. Reagent blanks, duplicated samples, and sample spike recoveries were all analyzed during each batch of analysis.

### 2.6. Cell viability

ExoFeR and ExoFeR^TF^ were treated at three time points (24, 48, 72 hours). for CDDP treatment ExoFeR and ExoFeR^TF^ were treated 1 hour prior to CDDP treatment. After each time point the cell were collected by trypsinization and made into a cell suspension. 0.4% Trypan Blue solution (Sigma, Cat. No.15250061) and cell suspension were made in 1:1 ratio mixture. The resulting mixture is loaded onto a hemocytometer, and cells are allowed to settle into counting chambers. Under a microscope, stained (blue) and unstained (clear) cells are counted in multiple squares. Cell viability is then calculated as the percentage of unstained cells relative to the total cell count. Unstained cells are considered viable, while stained cells indicate non-viability.

### 2.7. PKH67 labelling

ExoFeR and ExoFeR^TF^ were labeled using a PKH67 green fluorescent labeling kit (Sigma, Cat. No. PKH67GL-1KT) in accordance with the manufacturer’s protocol. Briefly, 6 μl of PKH67 were combined with 1 ml of Diluent C in microfuge tubes for each sample, as well as an additional tube for a media-only control. The resulting dye/diluent mixture was added to the ultracentrifugation tube for each sample, followed by gentle pipetting for 30 seconds. To stop the reaction, 2 ml of 10% BSA in PBS were introduced, and the volume was adjusted to 8.5 ml with serum-free media. A 0.971 M sucrose solution was prepared from a 2.5 M sucrose stock solution, and 1.5 ml of this solution was meticulously added to the bottom of the tube, creating a sucrose cushion beneath the EV/PKH67 solution. Following centrifugation at 110,000 × g for 2 hours at 4 °C, the samples were in the pellet, with most of the excess dye remaining in the interface layer. The media and interface layer were aspirated, and the EV pellet was gently resuspended through pipetting.

### 2.8. Western blotting

The cell lysis process involved the use of RIPA buffer (Sigma, Cat. No. R0278-500ML) combined with sonication. Protein concentrations in the cell lysate were determined through the application of the Pierce BCA Protein Assay Kit (Thermo Fisher Scientific, Cat. No. 23225). Subsequently, the cell lysate samples were subjected to a 10-minute heat treatment at 95°C with a dye (4x Laemmli Sample Buffer, BioRad, Cat. No. 1610747). Following this, proteins were separated through SDS-PAGE and then transferred onto polyvinylidene fluoride (PVDF) membranes from Sigma-Aldrich (Cat. No. IPVH00010). After blocking with a 3% BSA solution in Tris-buffered saline with 0.1% Tween-20 (TBST), the membranes underwent probing with specific primary antibodies as listed below in Table 1. Subsequently, HRP-conjugated secondary antibodies (Jackson Immuno Research, Cat. No. 111-035-144 and 115-035-146) were applied. Blots were developed using a chemiluminescence kit (BioRad, Irvine, Cat. No. 1705061) and captured using the Gbox mini system by Syngene in CA, USA.

**Table 1:** List of antibodies used in the study.

| No. | Antibody Name | Catalogue No. | Dilution | Company |
| --- | --- | --- | --- | --- |
| 1 | GPX-4 | sc-166570 | 1:1000 | Santa Cruz |
| 2 | xCT/SLC7A11 | 12691S | 1:2000 | Cell Signaling Technologies |
| 3 | Nrf2 | sc-365949 | 1:1000 | Santa Cruz |
| 4 | HO-1 | 70081S | 1:2000 | Cell Signaling Technologies |
| 5 | KEAP1 | sc-365626 | 1:1000 | Santa Cruz |
| 6 | Lamin a/c | sc-376248 | 1:1000 | Santa Cruz |
| 7 | Cytochrome c | sc-13156 | 1:1000 | Santa Cruz |
| 8 | GRP94 | ab210960 | 1:1000 | Abcam |
| 9 | β-actin | A2228 | 1:10000 | Sigma |

### 2.9. Zeta potential measurements

To assess the surface charge of particles, a laser doppler microelectrophoresis technique utilizing non-invasive backscatter technology was employed. Zeta potential, representing the surface charge at slipping planes, was determined using a Zetasizer (Nano ZS from Malvern Panalytical, Malvern, UK) with a 633 nm He–Ne laser source for electrophoresis light scattering (ELS). In a nutshell, a solution of EVs (200 µg), IONP (40 µg), ExoFeR (comprising 200 µg of EV and 40 µg of IONP) in water was prepared, and 800 μl of this solution underwent sonication (10 seconds), vortex mixing (5 seconds), and was then introduced into a DTS-1070 Zeta-measurement cell. For the sample parameters, specifications included the material designated as “Iron,” the Debye-Huckel approximation, the solvent indicated as “water,” and a temperature equilibrated to 25 °C. All measurements were conducted in triplicates, with each measurement consisting of 10 runs. Data analysis was executed using the Zetasizer software suite (ver 7.13), and the zeta potential values were exported to MS-Excel for plotting.

### 2.10. Cellular and mitochondrial ferrous iron detection

FerroOrange and Mito-FerroGreen probes from Dojindo, Prussian blue staining (Iron, Gomori Prussian Blue Stain Kit, Newcomer Supply) was employed to evaluate intracellular ferrous iron levels, as per manufacturer’s instructions (Fujimaki et al., 2019; Hirayama et al., 2018). A549 cells were cultured, and after treatment with IONP and ExoFeR, iron levels were assessed using subsequently outlined staining techniques.

#### 2.10.1. Prussian blue staining for intracellular ferrous iron levels

Prussian blue staining was employed to assess intracellular ferrous iron levels. Following hydration in 100% and 95% ethyl alcohols and thorough washing with distilled water, a fresh Ferrocyanide Working Solution was prepared using Hydrochloric Acid 20% (Aqueous) and Potassium Ferrocyanide 10% (Aqueous). After a 20-minute incubation, cells were rinsed, treated with Nuclear Fast Red Stain (Kernechtrot) in Solution C for 5 minutes, rinsed again, and dehydrated in 95% and 100% ethyl alcohol twice. The samples were then subjected to microscopic examination (photomicroscope, Olympus, CX 44).

#### 2.10.2. Cellular ferrous iron detection with FerroOrange

For cellular ferrous iron detection, cells were cultured on 6-well plates, treated with ExoFeR, washed thrice with PBS, and incubated with 1 μM FerroOrange for 30 minutes at 37 °C, 5% CO2. Subsequently, they were observed under a fluorescent microscope (Keyence microscope, BZ-X810).

#### 2.10.3. Mitochondrial iron assessment using Mito-FerroGreen

To assess mitochondrial iron, cells were washed with PBS, incubated with 5 μM Mito-FerroGreen for 30 min at 37 °C, 5% CO2, and then washed thrice with PBS before fluorescent microscopic examination (Keyence microscope, BZ-X810).

### 2.11. Statistical analysis

The study results were analyzed using statistical software called GraphPad Prism 9.0. We summarized continuous outcome variables using mean values and standard deviations. To compare multiple pairs, we used a two-way ANOVA model along with Dunnett’s test. Results with adjusted p-values below 0.05 were considered statistically significant. (* P < 0.05, ** P < 0.01, *** P <0.005, **** P <0.001)

## 3. Results

### 3.1. ExoFeR, a formulated combination of iron oxide nanoparticles and EVs, induces ferroptosis in lung cancer cells

A reduction in cell viability was observed when a complex of only EV-IONP, having no treatment modality involved was added to lung cancer (A549) cells (data not shown). We speculated these non-canonical cell deaths were due to ‘Ferroptosis’: a newly discovered form regulated cell death (RGD) that is mechanistically different from apoptosis and is induced by Iron (Fe^+2^ ions) known as Labile Iron Pools (LIPs). To further investigate this phenomenon two concentrations of iron oxide nanoparticles (IONP), were (20 µg and 40 µg) combined to 200 µg total protein weight equivalent of EVs and administered to A549 cells. The cell viability assay with this complex revealed that 20 and 40 µg IONP with EVs resulted in approximately 68.3 and 48.3% viable cells at 48 hours (Fig. 1a) whereas IONP alone treatment showed about 77.7 and 70% cell viability of 20 and 40 µg. Based on this data, we proceeded to formulate 40 µg of IONP conjugated with EVs all future studies.

**Figure 1.**
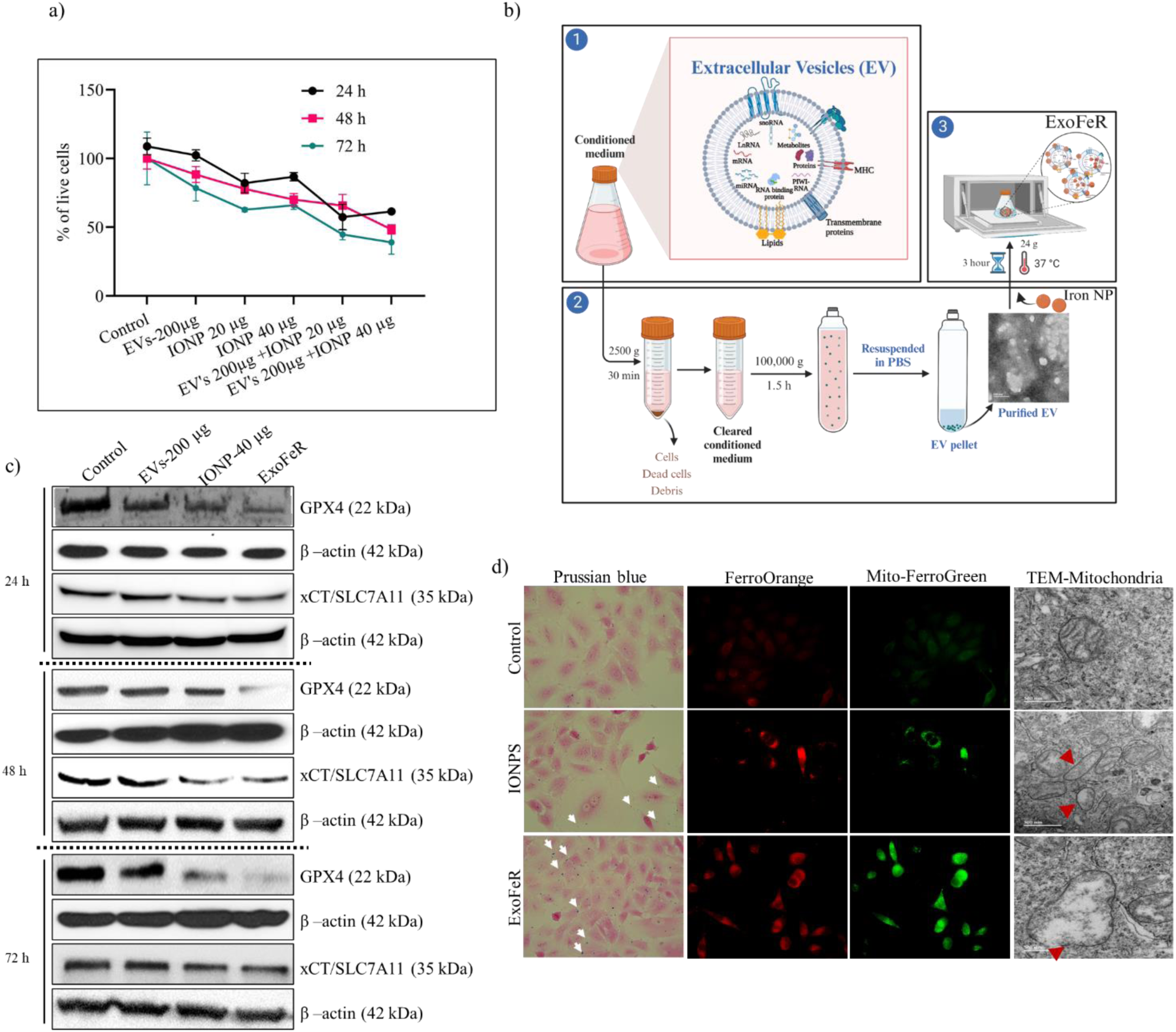
Comprehensive analysis of ExoFeR treatment effects in A549 cells. (**a**) Quantification of live cells expressed as a percentage after treatment with different concentrations of IONPS and EVs combined with IONPS using trypan blue staining (**b**) Schematic representation detailing the steps involved in the purification of EVs and the preparation of ExoFeR. (**c**) Western blot results depicting the expression levels of ferroptosis markers (GPX-4, XCT/SLC7A11) at three time points (24, 48, 72 hours) following treatment with ExoFeR and IONPS. (**d**) Visualization of ferric iron (Fe^3+^) and ferrous iron (Fe^2+^) in A549 cells using Prussian Blue. Intracellular free ferrous iron is detected by FerroOrange, and mitochondrial free ferrous iron is identified by Mito-FerroGreen. Mitochondrial morphology is assessed through transmission electron microscopy (TEM) imaging.

We coined term ‘ExoFeR’ for the formulation because of its composition that consisted of EVs (Exo) a natural nano carriers coupled with IONPs a known ferroptosis inducer (FeR) (Fig. 1b). The classic pathways of induction of ferroptosis involve the inhibition of the upstream regulator system xCT/SLC7A11 (an amino acid antiporter) or the downstream effector glutathione peroxidase 4 (GPX4) of GSH (D. Tang & Kroemer, 2020). We examined both xCT/SLC7A11 and GPX4 expression levels in response to the treatment of IONP alone and ExoFeR. Western blot analysis exhibited a time-dependent downregulation of xCT/SLC7A11 and GPX4 with ExoFeR treatment (Fig 1c), indicating that ExoFeR induces ferroptosis in the treated cancer cells.

Further, since the accumulation of iron is linked to increased ROS production that plays a crucial role in ferroptosis; therefore, it is speculated that both cytosolic and mitochondrial iron are essential contributors to the ferroptotic process (Chen et al., 2023; S. Zhang et al., 2022). Examining the increased levels of intracellular or mitochondrial ferrous iron (Fe^2+^) in the putative ferroptotic cells after the treatment with ExoFeR would further prove the ferroptosis-inducing capability of the system. Prussian blue (Lu et al., 2020), FerroOrange (Tomita et al., 2019), and Mito-FerroGreen (F. Yu et al., 2022) staining revealed a significant increase in the deposition of free iron in cells and mitochondria in the ExoFeR-treated group relative to the untreated control (Fig 1d).

A structural hallmark of ferroptosis is morphological changes in mitochondria, including increased membrane density and reduced or disappearance of mitochondrial cristae (M. Gao et al., 2019). TEM analysis of ExoFeR-treated cells demonstrated the almost complete disappearance of cristae (Fig 1d). Thus, the downregulation of ferroptosis markers (GPX-4 and xCT/SLC7A11), the presence of free iron in cells and mitochondria, and morphological changes in mitochondria collectively support the assertion that ExoFeR can induce ferroptosis in lung cancer cells.

### 3.2. EVs Characterization

In adherence to the MISEV criteria (Welsh et al., 2024), we performed thorough characterizations of individual EVs. TEM and NTA of EVs confirmed typical EV morphological characteristics of double membrane circular shape having an average diameter size ranging from 150.6 nm to 190 nm (Fig. 2a). The MISEV guidelines do not suggest using widely applicable molecular markers. Instead, they recommend looking at at-least one transmembrane or cytosolic protein to show the unique features of EV and differentiate them from cell debris. Even though the guidelines do not provide specific recommendations for universal negative markers, they stress the importance of carefully removing contaminants and major components of non-EV structures. This step is crucial for thoroughly checking the purity of the EV formulation. With this in mind, we analyzed ESCRT (endosomal sorting complex required for transport) proteins, such as ALIX and TSG101 that are involved in EV cargo sorting (Raiborg & Stenmark, 2009) and biogenesis. Similarly, Flotillin-1 (a lipid raft protein) and tetraspanins member protein CD9 are frequently used to detect and confirm the presence of EVs. Western blot analysis showed that the EVs isolated from MRC-9 exhibited enrichment of EV-marker proteins *viz.* ALIX, TSG-101, flotillin-1, CD9. Further, to negate the possibility of contamination from other subcellular components in the EV preparations the western blots showed absence of endoplasmic reticulum marker (Grp94), nucleus marker (Lamin A/C), and mitochondrial protein marker (cytochrome c) in comparison to the cell lysates (Fig. 2b). Following MISEV guidelines, we characterized individual EVs (diameter: 150.6–190 nm) using TEM and NTA. Utilizing transmembrane or cytosolic proteins as recommended, including ALIX, TSG101, Flotillin-1, and CD9, we confirmed EV presence in MRC-9 isolates *via* Western blot. Absence of endoplasmic reticulum, nucleus, and mitochondrial markers in comparison to cell lysates ensured EV purity.

**Figure 2.**
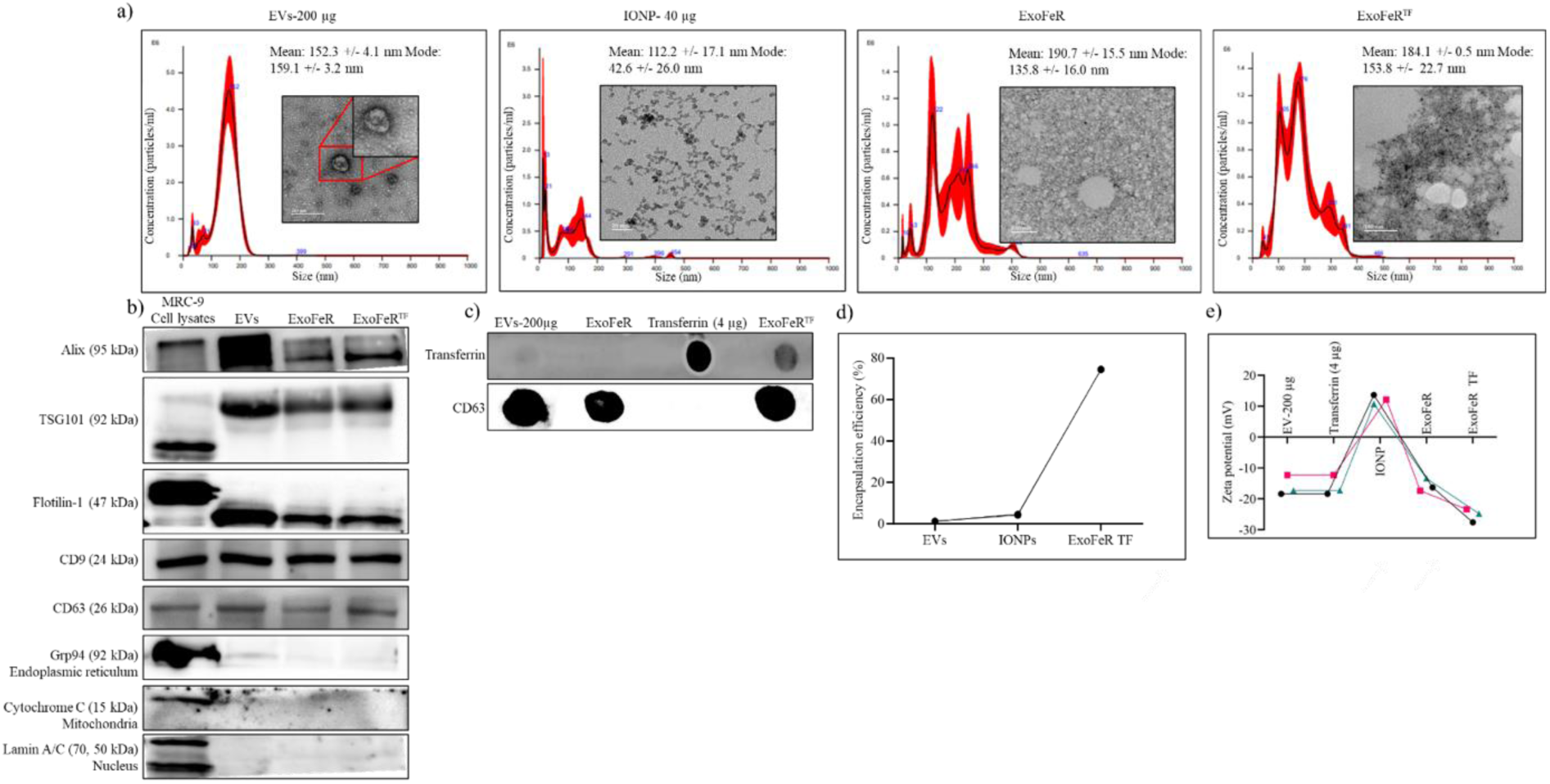
Multi-modal analysis of ExoFeR and ExoFeR^TF^: characterization, and intracellular dynamics. (**a**) Characterization of EV, IONP, ExoFeR, and ExoFeR^TF^ using NTA and TEM. **(b)** Western blot analysis was conducted to assess membrane and intraluminal proteins characteristic of EVs. The markers examined included ALIX, TSG-101, flotillin-1, CD9, and the endoplasmic reticulum marker (Grp94), nucleus marker (Lamin A/C), and mitochondrial protein marker (cytochrome c). (**c**) Confirmation of transferrin presence in ExoFeR^TF^ by Dot blot. **(d)** ICP-MS analysis was performed to evaluate the efficiency of complexation between IONPs and EVs in ExoFeR^TF^. **©** Zeta potential (ZP) analysis was utilized to examine the surface potential of EVs, assessing both surface charge and colloidal stability of ExoFeR and ExoFeR^TF^.

### 3.3. Transferrin-conjugated (ExoFeR^TF^) outperforms ExoFeR, inducing higher cell death, with improved uptake

Studies indicate elevated TfR expression in proliferating normal and cancer cells compared to resting cells, likely due to increased iron needs for DNA synthesis. TfR has long been a target for pharmacological intervention due to its differential expression in normal and malignant tissues particularly in lung cancer tissues (Kukulj et al., 2010a; H. Li & Qian, 2002). The increase in TfR in lung cancer cells suggests that it could increase iron intake by enhancing the effects of the TF and TfR. Thus, we combined TF with ExoFeR to increase its efficacy and for a more targeted approach.

Dot blot assay indicated TF binding with ExoFeR^TF^ and not in ExoFeR (Fig. 2c). Inductively coupled plasma mass spectrometry (ICP-MS) is a method that was employed to estimate the encapsulation efficiency with iron concentration in ExoFeR^TF^. The analysis of the supernatant and pellet revealed approximately 30 µg of iron concentration in the ExoFeR^TF^ pellet, representing about 75% complexation efficiency compared to the initially supplied 40 µg of iron (Fig. 2d).

EVs play a vital role in intercellular communication and typically carry a negative surface charge. Zeta potential (ZP) analysis is commonly used to measure their surface potential, indicating surface charge and colloidal stability. Our analysis showed that EVs, ExoFeR, and ExoFeR^TF^ all exhibited a negative charge within the acceptable range of variation (Helwa et al., 2017; Kukulj et al., 2010b; Y. Wang et al., 2015), mostly below -10 mV, ensuring sample stability in dispersion (Fig. 2e).

The characteristics of EVs remained the same in both the ExoFeR and ExoFeR^TF^ complexes, confirmed by consistent protein patterns (ALIX, TSG-101, flotillin-1, CD9) in western blot analysis. Moreover, absence of endoplasmic reticulum, nucleus, and mitochondrial markers in western blots compared to cell lysates ensured the absence of contamination (Grp94, Lamin A/C, cytochrome c). TEM and NTA showed typical EV shapes and an average size of about 190 nm, though NTA revealed an increase in heterogeneity EVs.

Further, cell viability analysis demonstrated increased cell death with ExoFeR^TF^ (55.7%) compared to ExoFeR (21.7%) and the control after 24 hours. After 48 hours, ExoFeR^TF^ exhibited 60.6 % cell death, while ExoFeR showed 47.65 % (Fig. 3a). We also assessed the MRC-9 lung fibroblast cell line, revealing approximately 20% cell death at 48 hours (Supplementary. Fig. 1a). This demonstrates that ExoFeR and ExoFeR^TF^ are non-toxic to the normal cells. Treatment with the EV inhibitor (GW4869) resulted in no change, indicating that the effect observed with ExoFeR treatment is not due to intrinsic effects from the EVs themselves (Supplementary. Fig. 2a, b). Western blot analysis of ferroptosis markers further supported the increased effectiveness of ExoFeR^TF^ (Fig. 3b). These results indicate combining TF with ExoFeR improved targeting and efficacy, as observed in cell viability assays and western blots.

**Figure 3.**
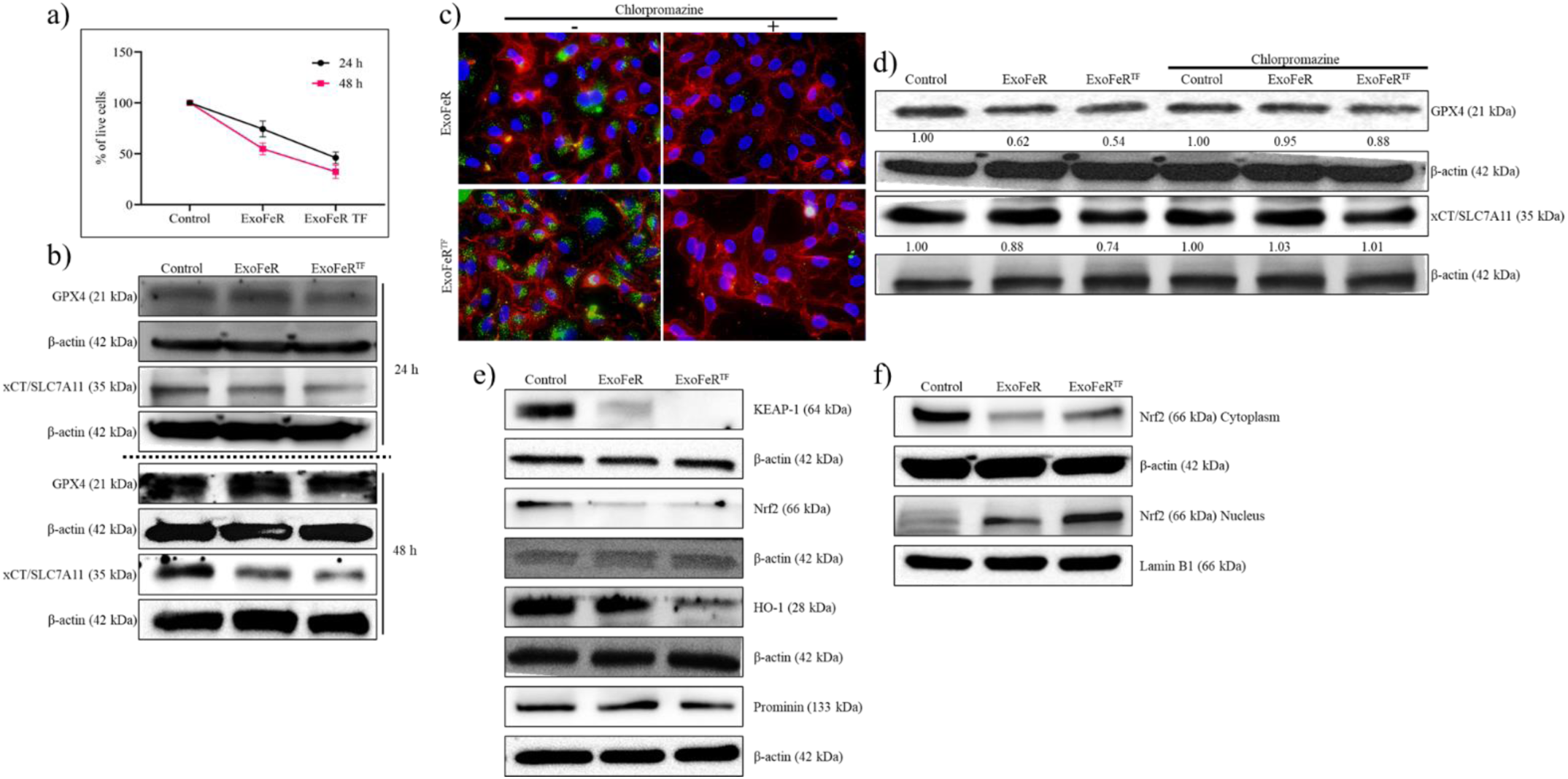
Comprehensive Investigation of ExoFeR and ExoFeR^TF^: Cell Viability, Ferroptosis Markers, Uptake Mechanisms, and Nrf2-Mediated Responses. (**a**) Quantification of cell viability via Trypan Blue staining after ExoFeR and ExoFeR^TF^ treatment. (**b**) Western blot analysis of ferroptosis markers (GPX-4, XCT/SLC7A11) at different time points (24, 48 h) following ExoFeR and ExoFeR^TF^ treatment. (**c**) Investigation of uptake and endocytic trafficking of ExoFeR and ExoFeR^TF^, with inhibition of clathrin-mediated endocytic pathways. (**d**) Western blot analysis of ferroptosis markers (GPX-4, XCT/SLC7A11) following ExoFeR and ExoFeR^TF^ treatment, with inhibition of clathrin-mediated endocytic pathways. (**e**) Western blot analysis of Nrf2 and downstream proteins **(f)** Nrf2 translocation from cytoplasm to nucleus after treatment of ExoFeR and ExoFeR^TF^.

### 3.3. ExoFeR and ExoFeR^TF^ undergo internalization through clathrin-dependent endocytosis

EV transport cargo and influence target cells through processes such as membrane fusion, endocytosis, and micropinocytosis. Understanding the endocytosis pathway of EV is vital for optimizing delivery efficiency, targeting strategies, and minimizing immune responses, ultimately enhancing their therapeutic potential (Kanada et al., 2015; Zomer et al., 2015). To identify the specific uptake pathway involved with ExoFeR^TF^, we employed inhibitors of different endocytosis pathways, including amiloride (blocking H^+^/Na^+^ and Na^+^/Ca^2+^ channels regulating calcium levels), omeprazole (a proton pump inhibitor), genistein (a tyrosine-kinase inhibitor), Dynasore (clathrin-dependent), and chlorpromazine (clathrin-mediated) (Supplementary Fig. 3a). Chlorpromazine efficiently inhibits clathrin-dependent endocytosis by promoting the assembly of clathrin and adaptor proteins, such as TF, on endosomal membranes, resulting in the depletion of clathrin from plasma membranes. Notably, Chlorpromazine significantly decreased ExoFeR^TF^ uptake, and western blot analysis of ferroptosis markers indicated that ExoFeR^TF^ follows clathrin-mediated endocytosis (Fig. 3c, d). These findings indicate that ExoFeR^TF^ internalization is through clathrin-mediated endocytosis.

### 3.4. Nrf2 translocation in ExoFeR^TF^-modulated ferroptosis

Nrf2 plays a crucial role in preventing ferroptosis by controlling iron levels, antioxidant defense enzymes, and regulating glutathione, thioredoxin, and NADPH. Ferroptosis is triggered by inhibiting XCT/SLC7A11 and GPX4, both of which are regulated by Nrf2. Nrf2 is also a transcription factor for XCT/SLC7A11 (Fan et al., 2017; Hayes & Dinkova-Kostova, 2014; Shin et al., 2018; Sun et al., 2016). Therefore, we investigated the impact of ExoFeR^TF^ treatment on the status of Nrf2 and downstream proteins. The expression of Nrf2 was decreased with ExoFeR^TF^ treatment (Fig. 3e).

Under normal conditions, Nrf2 forms a complex with Kelch-like epichlorohydrin-associated protein 1 (Keap1). In oxidative stress, Nrf2 dissociates, moves to the nucleus, activates antioxidant response elements, and enhances the genes like xCT/SLC7A11, reinforcing system XCT’s antioxidant effect. ExoFeR^TF^ treatment impedes Nrf2 translocation into the nucleus, suggesting ExoFeR^TF^ inhibits Nrf2 and induces ferroptosis by limiting its nuclear translocation (Fig. 3f). Thus, ExoFeR^TF^ treatment was found to decrease Nrf2 expression, inhibiting its nuclear translocation and subsequent activation of antioxidant response elements, including xCT/SLC7A11. This disruption of Nrf2 signaling contributes to ferroptosis induction by ExoFeR^TF^, underscoring its potential as a therapeutic strategy.

### 3.5. Sensitization of A549, HCC827, and cisplatin-resistant tumoroids to cisplatin by ExoFeR and ExoFeR^TF^

Ferroptosis is considered a potential mechanism to overcome resistance to cisplatin (CDDP) (Golbashirzadeh et al., 2023). CDDP-resistant cells often develop resistance mechanisms against apoptosis, a common cell death pathway induced by conventional chemotherapeutics. Ferroptosis offers an alternative pathway that is not easily evaded (Z. Tang et al., 2021) by resistant cells, providing a novel approach to overcome cisplatin resistance and enhance the effectiveness of treatment. Building on our previous results, which established that ExoFeR^TF^ induces ferroptosis, we next aimed to explore whether induction of ferroptosis makes any changes in CDDP sensitivity in cancer cells. We also investigated whether combining ExoFeR^TF^ with CDDP could improve effectiveness and make cells more responsive to CDDP. We chose A549 and HCC827 cell lines because they respond differently to CDDP, with A549 having a high IC50 (20 μM) value and HCC827 (10 μM) having a low IC50 value. Additionally, we also incorporated tumoroids derived from NSCLC patients (Table 2) which showed resistance to CDDP. The results demonstrated a heightened sensitivity to CDDP in the presence of ExoFeR and ExoFeR^TF^ in all cell lines tested. Cell viability assays revealed a significant reduction in the survival rates of A549, HCC827, and CDDP-resistant patient-derived tumor organoids (PDTOs) (MU387, MU369, MU383) when treated with the combination of ExoFeR or ExoFeR^TF^ with CDDP. The cell viability of A549 treated with ExoFeR^TF^ and CDDP was about 10 % at 72 h whereas HCC827 showed 17 %. The resistant PDTOs showed less than 5 % viability (Fig 4a, b, c). Furthermore, ferroptosis markers (GPX-4, xCT/SLC7A11) were studied to elucidate the mechanistic aspects of the sensitization process. We also examined the impact of ExoFeR^TF^ on commercially available and established CDDP resistant ovarian cancer cells, namely A2780 and A2780-cis resistant cells. The results showed a reduction in cell viability and the inhibition of ferroptosis markers (GPX-4, xCT/SLC7A11) with ExoFeR^TF^ treatment, emphasizing the fact that ExoFeR^TF^ effect is consistent (Supplementary Fig. 6a, b).

**Figure 4.**
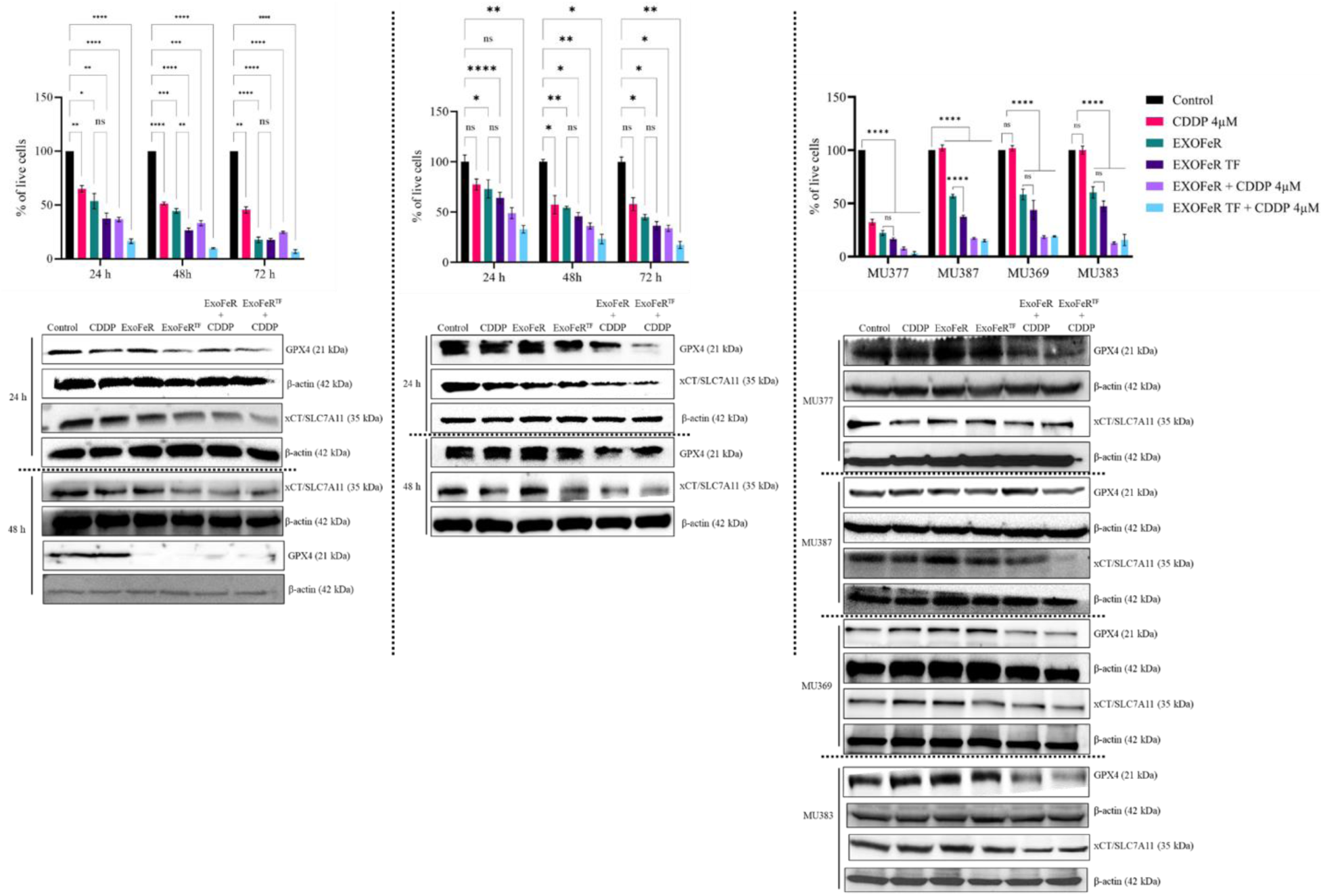
ExoFeR and ExoFeRTF sensitize CDDP treatment. Quantification of viable cells with the treatment of ExoFeR and ExoFeR^TF^ with CDDP and western blot analysis of ferroptosis markers (GPX-4, XCT/SLC7A11) **(a)** A549 **(b)** HCC827 **(c)** PDTOs.

**Figure 5.**
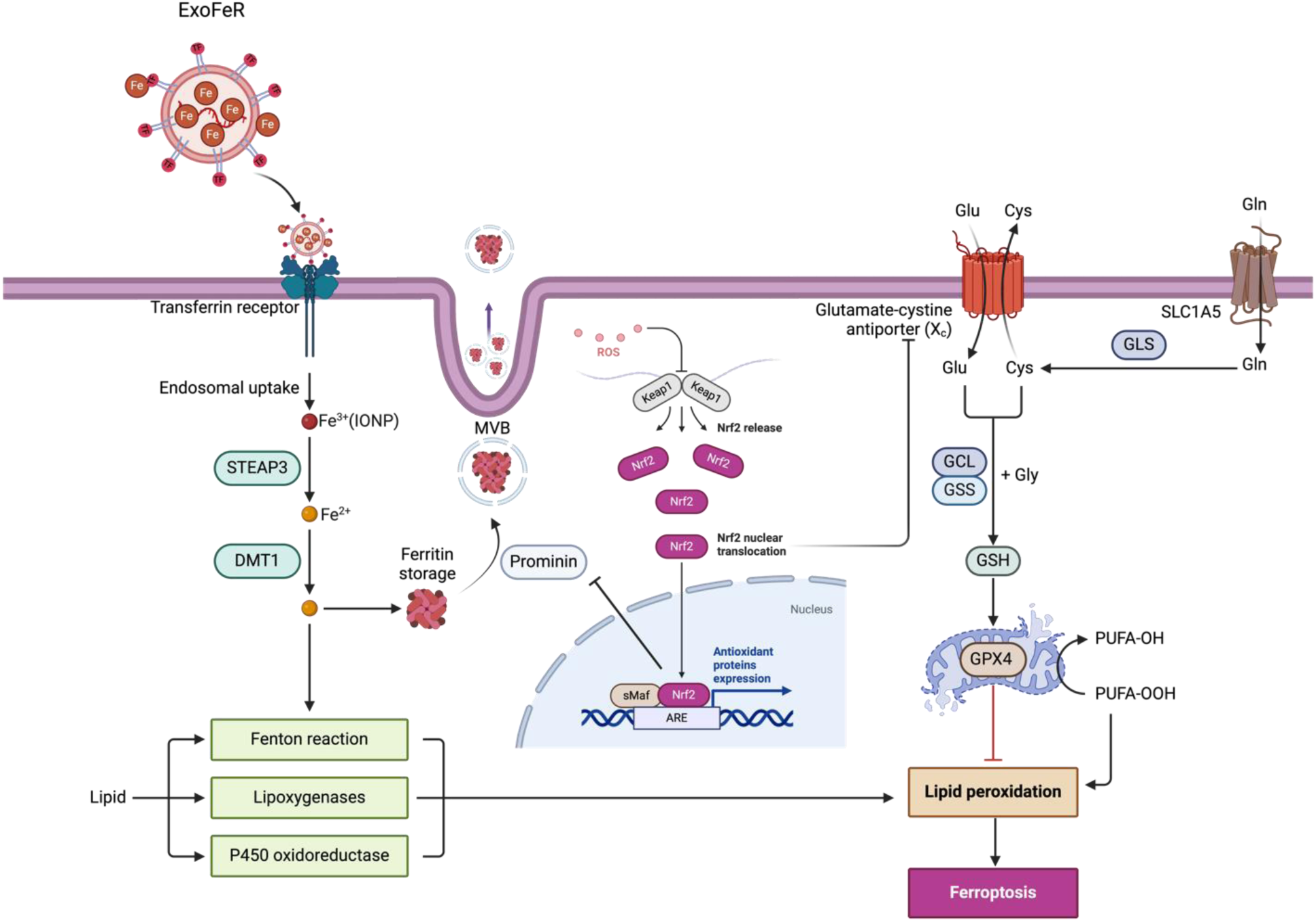
Schematic diagram of ExoFeR^TF^ uptake and functional mechanism.

**Table 2:** Clinicopathological information, PDO treatment responses and recurrence status of the surgical NSCLC patients enrolled (N=4). **Abbreviations:** NSCLC: non-small cell lung cancer, LUAC: lung adenocarcinoma, LUSC: lung squamous cell carcinoma, AJCC: American Joint Committee on Cancer, PDTO: patient-derived tumor organoid; N/A: not applicable. *All patients received adjuvant therapy targeting micrometastatic disease. In the adjuvant setting, drug responses could not be evaluated as the tumors were surgically removed at the time of drug administration (i.e., there was no radiographically measurable disease left for treatment response evaluation).

| *Patient ID* | *Age/ Gender* | *NSCLC histologic subtype* | *Smoking history (pack years; former/ current)* | *Tumor stage (AJCC)* | *PDTO response to carboplatin + paclitaxel* | *Adjuvant treatment after surgery* | *Response to adjuvant treatment\** | *Months of observation after surgery* | *Recurrence /Death* | *Months until recurrence (site)* | *Treatment for recurrence* |
| --- | --- | --- | --- | --- | --- | --- | --- | --- | --- | --- | --- |
| MU369 | 71/M | LUSC | 50; current | IA | **Resistant** | None | N/A* | 32 | No/No | N/A | N/A |
| MU377 | 68/F | LUSC | 70; former | IA | **Sensitive** | None | N/A* | 31 | No/No | N/A | N/A |
| MU383 | 64/M | LUAC | 40; former | IIIA | **Resistant** | Carboplatin + Paclitaxel | **Resistant (Recurrence during treatment)** | 15 | **Yes/Yes** | 2 (*Lymph nodes*)<br>10 (*Brain*) | Atezolizumab<br>Radiation |
| MU387 | 65/M | LUSC | 40; current | IA | **Resistant** | None | N/A* | 30 | No/No | N/A | N/A |

In conclusion, our research highlights the promising role of ExoFeR and ExoFeR^TF^ in enhancing the effectiveness of CDDP across various cell lines. This includes cells that developed resistance due to continuous cisplatin treatment (A2780-cis), inherent resistance (A549), and sensitive cells (HCC827), along with cisplatin-resistant PDTOs.

## 4. Discussion

The functional significance of ferroptosis as a non-canonical regulated cell death mechanism is well recognized for its benefit in cancer therapy, such as by overcoming cisplatin resistance in cancer cells (Cheng et al., 2021; D. Fu et al., 2021; R. Fu et al., 2024). Iron oxide nanoparticles (IONPs) can naturally trigger ferroptosis; however, direct infusion in the body’s complex blood flow system tends to accumulate IONPs prominently in the liver and spleen (Hofmann et al., 2014) resulting in compromised bioavailability to tumor sites. Studies have identified that IONPs uptake by cells occurs through endocytosis (Liu et al., 2019; Luther et al., 2013; Petters et al., 2014) and thus suitable to enhance bioavailability by encapsulating IONPs in a polymeric nanoparticle. Though this strategy is promising, it remained unfeasible for clinical application due to its synthetic nature and absence of tumor targeting (J.-H. Kim et al., 2017). EVs share notable structural similarities with lipid-based polymeric nanoparticles and have been extensively explored as therapeutic delivery vehicles (Le Saux et al., 2021; Mitchell et al., 2021; Rodríguez & Vader, 2022). EVs offer several advantages over conventional lipid-based nanoparticles as they are natural in origin without any immunogenic effect, have better bioavailability and *de novo* organotropism, and can be functionalized to confer tumor-targeting abilities (Rosso & Cauda, 2023; Srivastava et al., 2022). In the present study, we successfully demonstrated the intrinsic induction of ferroptosis in lung cancer cells and PDTOs by delivering IONPs through EVs and termed this novel system as ExoFeR. The simplistic formulation of ExoFeR and the organic ability to induce ferroptosis without any significant off-target toxicities place it as a promising choice for clinical applications.

To enhance efficiency, we explored the unique features of TF receptors, especially for improving tumor-targeted delivery. Several studies have shown higher expression of glycoprotein TfR (CD71) on NSCLC cell surfaces and its involvement in intercellular iron transport and cell growth (Kukulj et al., 2010a; Parvathaneni et al., 2021; Upadhyay et al., 2019; Z. Wang et al., 2024). This suggests an opportunity to increase iron absorption by strengthening the impact of TF and transferrin receptor (TfR). ExoFeR^TF^, enriched with iron, naturally enters cells through TfRs, surpassing the usual endocytosis barriers leading to enhanced ferroptosis in cancer cells. The lipophilic dye PKH67-labeled ExoFeR^TF^ showed higher cellular uptake compared to ExoFeR (Fig. 2c), highlighting the effectiveness of TF in promoting iron absorption and emphasizing crucial contribution of tumor targeting by TF for boosting the therapeutic capabilities of ExoFeR^TF^. We employed an EV inhibitor (GW4869) to ensure that the observed ferroptosis induction is due to ExoFeR and not inherent to EVs or any bystander entities generated by the recipient cells. The induction of cell death by ExoFeR^TF^ in the presence of the EV inhibitor conferred that the effects are indeed caused by ExoFeR^TF^. We also verified the nonapoptotic nature of ferroptosis by using an inhibitor for apoptosis (Z-VAD-FMK) and demonstrated the exclusivity of cell death or RGD by ExoFeRTF-induced ferroptosis. In contrast, a ferroptosis inhibitor (ferrostatin-1) decreased cell death with no impact on ferroptosis marker proteins. Through several rigorous studies we have proved that ExoFeR^TF^ is an and efficient and viable system to promote ferroptosis in lung cancer cells. A recent study also conferred our premise and demonstrated ferroptosis induction using iron nanoparticles and EV (exosomes) complex (ESIONPs@EXO) with an iron concentration of 54.37 ng per 1 μg of EVs [36]. In contrast, our complex induced ferroptosis with only 30.57 ng of iron per 200 μg of EVs, highlighting its significantly higher efficiency.

Nrf2 exerts a positive influence on xCT/SLC7A11, a key transporter responsible for importing cystine and exporting glutamate (Lewerenz et al., 2013). Another component in this regulatory network is SLC1A5 (solute-linked carrier family A1 member 5) that is activated by Nrf2, playing a pivotal role in glutamine uptake, thereby contributing to the biosynthesis of glutathione (GSH) (He et al., 2020). KEAP1, acting as the endogenous inhibitor of Nrf2, binds to Nrf2 and redirects it for proteasomal degradation. The KEAP1/Nrf2 interaction is crucial in maintaining Nrf2 at a basal level. Clinically relevant KEAP1 mutations disrupt this interaction, leading to elevated Nrf2 levels and drug resistance. Notably, treatment with ExoFeR and ExoFeR^TF^ in KEAP1 mutant lung cancer cells A549 cells resulted in the downregulation of Nrf2 proteins. A recent study by Aboulkassim et al. (Aboulkassim et al., 2023) corroborates these findings, demonstrating that R16, an Nrf2 inhibitor, sensitized KEAP1-mutant lung cancer cells to chemotherapy. While the precise mechanism by which ExoFeR^TF^ directly impedes the expression of Nrf2 remains to be fully elucidated, existing evidence strongly suggests the noteworthy potential of ExoFeR and ExoFeR^TF^ in modulating Nrf2 and hence regulate the GSH biosynthesis in cancer cells.

To gain the therapeutic benefit against cancer cells, induction of ferroptosis by IONPs requires enrichment of Fe (II) ions, which instigates fenton reaction within the acidic tumor microenvironment. This catalytic process converts H2O2 into highly toxic •OH , elevating cellular oxidative stress or ROS species causing damage to mitochondria and the cell membrane by lipid peroxidation (Zhu et al., 2022). However, generating sufficient •OH for induction of ferroptosis is challenging in cancer cells, as it is constrained by the limited availability of H2O2 required for the fenton reaction. To address this limitation, several studies demonstrated that an effective anti-tumor outcome can be achieved through an intracellular cascade of reaction mediated by the coordinated transport of metal catalysts that augments H2O2 concentration, and thus enhances the substantial production of •OH resulting in efficiently inducing cancer ferroptosis (Lin et al., 2019; Feng et al., 2018; Ma et al., 2017). Since cisplatin (CDDP) contributes indirectly to H2O2 production that instigates an intracellular cascade reaction, generating adequate •OH for iron-induced cell death treatment further accelerates the fenton reaction (Z. Gao et al., 2020; Hu et al., 2022; Shen et al., 2018). This mechanistic insight clarifies why ExoFeR^TF^ exhibits increased ferroptosis in combination with CDDP treatment. Interestingly, while CDDP itself inhibits GPX-4 and xCT/SLC7A11 in CDDP-sensitive lung cancer PDTOs (MU377), this effect was not observed in resistant PDTOs MU387, MU369, and MU383, suggesting a nuanced interplay between cisplatin sensitivity and ferroptotic pathways (Fig.4c) (Supplementary Fig 5).

In conclusion, this study highlights the transformative potential of EV-mediated therapies, particularly in combating cisplatin resistance and inducing ferroptosis for enhanced lung cancer treatment. The research illuminates the pivotal role of EVs in developing novel platforms for efficiently delivering IONPs to tumor cells and instigating a ferroptosis-mediated reversal of resistance against chemotherapeutic drug cisplatin. The developed vehicles ExoFeR and tumor-targeted transferrin-conjugated ExoFeR (ExoFeR^TF^), overcome several barriers of high precision cellular uptake offering superior efficacy in inducing ferroptosis in cancer cells. Further, unraveling the intricate cascade reactions involving ExoFeR^TF^ and CDDP holds promise for novel therapeutic strategies for overcoming chemotherapeutic drug resistance in cancer cells.

This study lays a foundation for a nuanced understanding of the interplay between ferroptosis induction, cisplatin sensitization, and Nrf2 modulation by ExoFeR^TF^, providing a valuable framework for future investigations in oncology therapeutics in lung cancer and beyond.

## Availability of data and materials

All data generated or analyzed during this study, if not included in this article and its supplementary information files, are available from the corresponding author on reasonable request.

## Ethics approval statement

All experiments conducted received approval from the University of Missouri School of Medicine’s Institutional Biosafety Committee (IBC) and Patients samples used in the study were obtained with written consent following the guidelines of the Institutional Review Board (IRB) of the University of Missouri (MU) through an approved protocol (IRB#: 2010166).

## Authors contribution

AP, RA, AS, JK: Conceived the idea and designed experiments. AP, YM, JK, AS: Developed PDTOs and performed ExoFeR efficacy assays. AP, RA, LX, HY: Characterized ExoFeR and ExoFer^TF^, performed and analyzed ICP-MS data. AP, RA, GM, SD, YM: performed and analyzed wet lab experiments. AP, RA, LX, HY, JK, AS: wrote and edited the manuscript. AS and JK: supervised the study.

## Acknowledgments

The authors thank Dr. Raghuraman Kannan and Dr. Dhananjay Suresh from the University of Missouri for providing access to the ZetaSizer and for their assistance with the measurement of zeta potential. The authors also acknowledge the Molecular Cytology Core, and to the Electron Microscopy (EM) Core at the University of Missouri, Columbia, Missouri, USA, for their assistance with confocal imaging and EM imaging respectively.

