## Supplementary figure for "Induction of Ferroptosis by an Amalgam of Extracellular Vesicles and Iron Oxide Nanoparticles Overcomes Cisplatin Resistance in Lung Cancer"

**Supplementary Figure 1.**

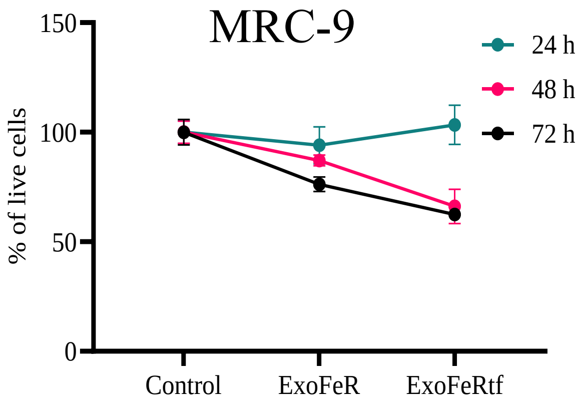


**Supplementary Figure 1. ExoFeR and ExoFeR$^{TF}$ treatment of MRC-9, a human lung fibroblast cell line.** The cell viability assay indicates that ExoFeR and ExoFeR$^{TF}$ treatment resulted in approximately 30% cell death, demonstrating that ExoFeR and ExoFeR$^{TF}$ are not toxic to the cells.

**Supplementary Figure 2.**

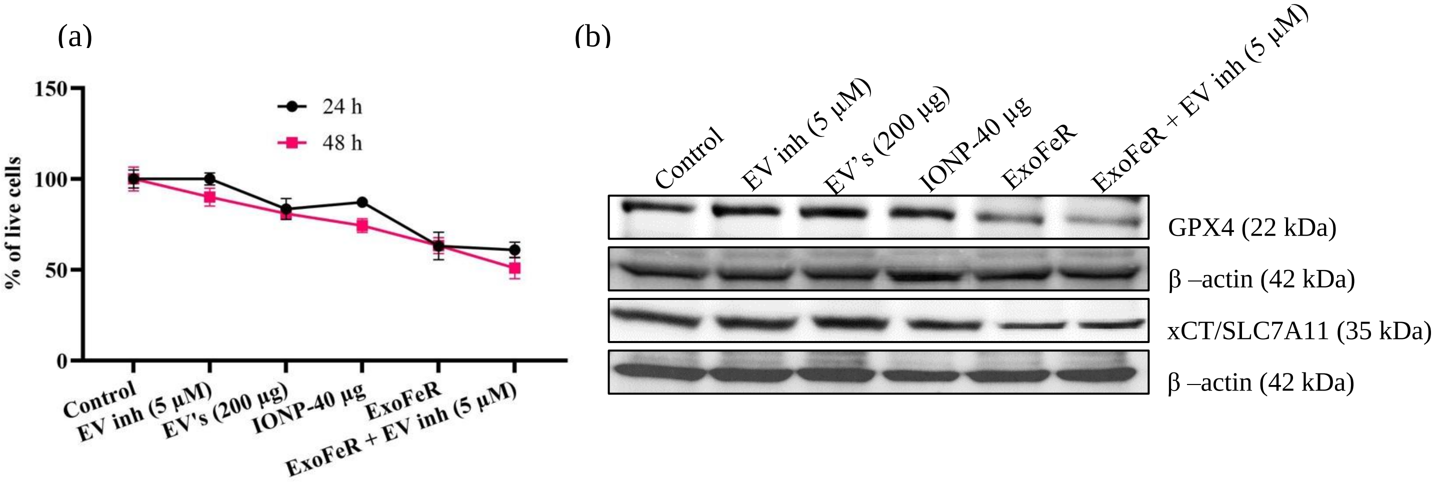


**Supplementary Figure 2. Treatment with an EV inhibitor with ExoFeR. (a)** A cell viability assay following EV inhibitor treatment alongside ExoFeR revealed approximately 62% cell death, indicating the effect is attributed to ExoFeR. **(b)** Western blot analysis of ferroptosis markers at 72 ho showed that the inhibition of GPX4 and xCT/SLC7A11 was not impeded by the EV inhibitor.

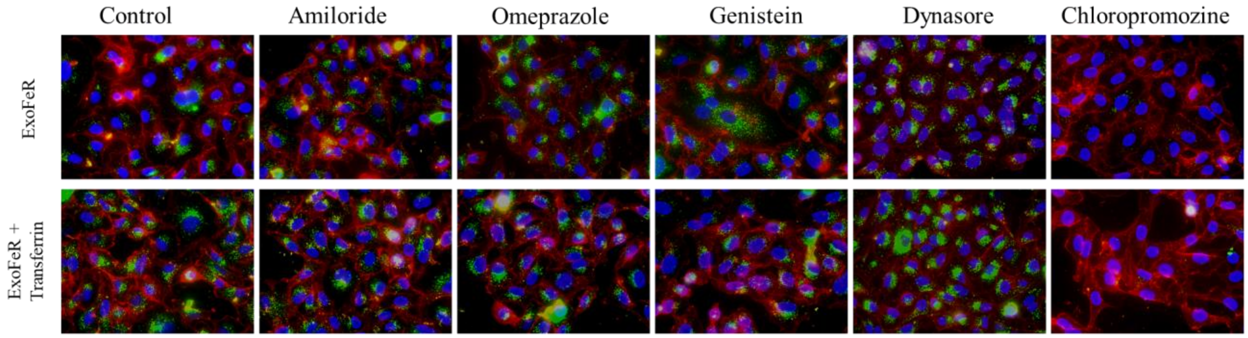


**Supplementary Figure 3. Analysis of internalization of ExoFeR and ExoFeR$^{TF}$.** Various inhibitors were used to determine the endocytosis internalization pathway of ExoFeR and ExoFeR$^{TF}$. Amiloride (blocks H+/Na+ and Na+/Ca2+ channels regulating calcium levels), omeprazole (a proton pump inhibitor), genistein (a tyrosine-kinase inhibitor), Dynasore (Clathrin-dependent), and chlorpromazine (clathrin-mediated) were tested. Among these, chlorpromazine was found to inhibit the internalization of ExoFeR and ExoFeR$^{TF}$.

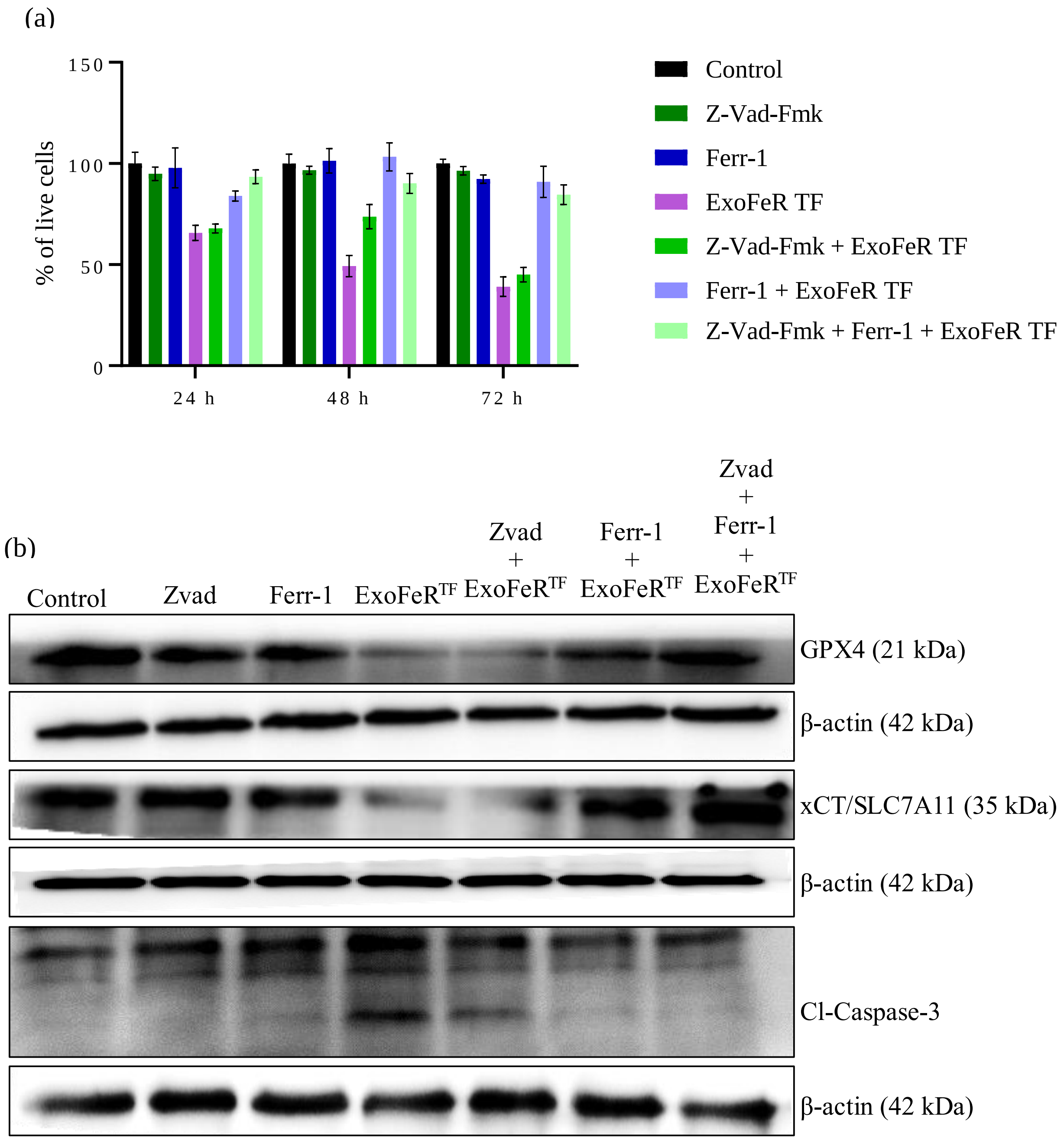


**Supplementary Figure 4. Pretreatment with Z-VAD-FMK, an apoptosis inhibitor, does not rescue cells from ExoFeR-induced cell death, whereas Ferrostatin-1, a ferroptosis inhibitor, does. (a)** Cell viability with Z-VAD-FMK and ExoFeR showed approximately 60% cell death, while Ferrostatin-1 with or without Z-VAD-FMK treatment resulted in about 10% cell death. **(b)**

Western blot analysis of ferroptosis markers and caspase-3 demonstrated that Z-VAD-FMK treatment did not affect GPX4 and xCT/SLC7A11 inhibition, whereas ferrostatin-1 did.

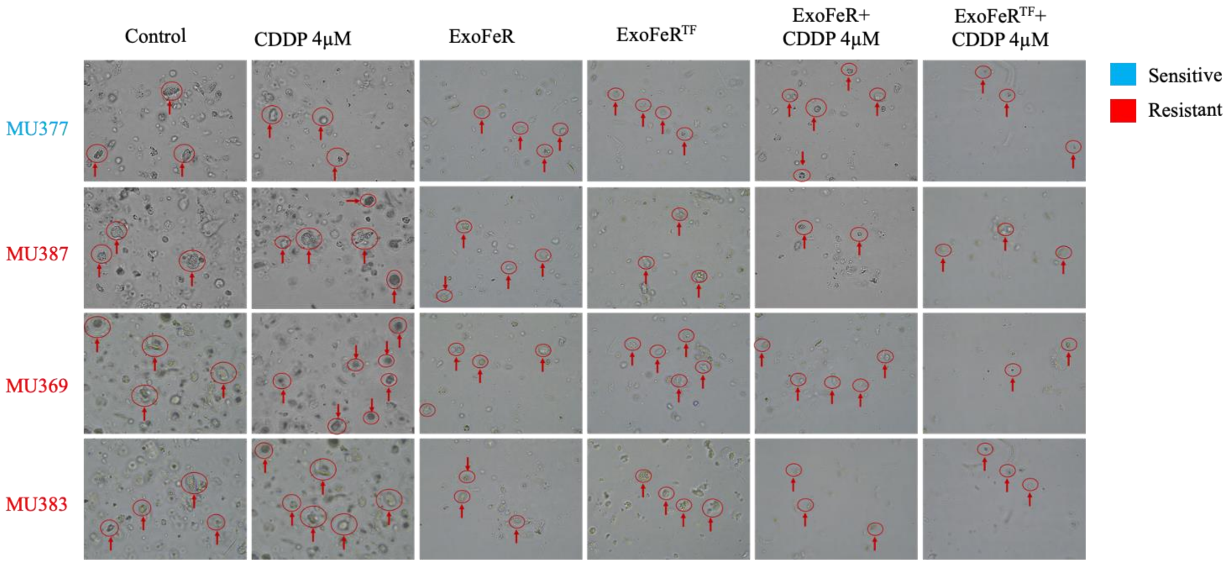


**Supplementary Figure 5. Treatment of ExoFeRTF and CDDP in patient-derived tumor organoids.** Both CDDP-sensitive (MU377) and resistant (MU387, MU369, MU383) tumor organoids were subjected to CDDP treatment with ExoFeR, resulting in fewer organoids compared to the untreated group. This indicates that ExoFeR reversed CDDP resistance in the organoids.

(a)

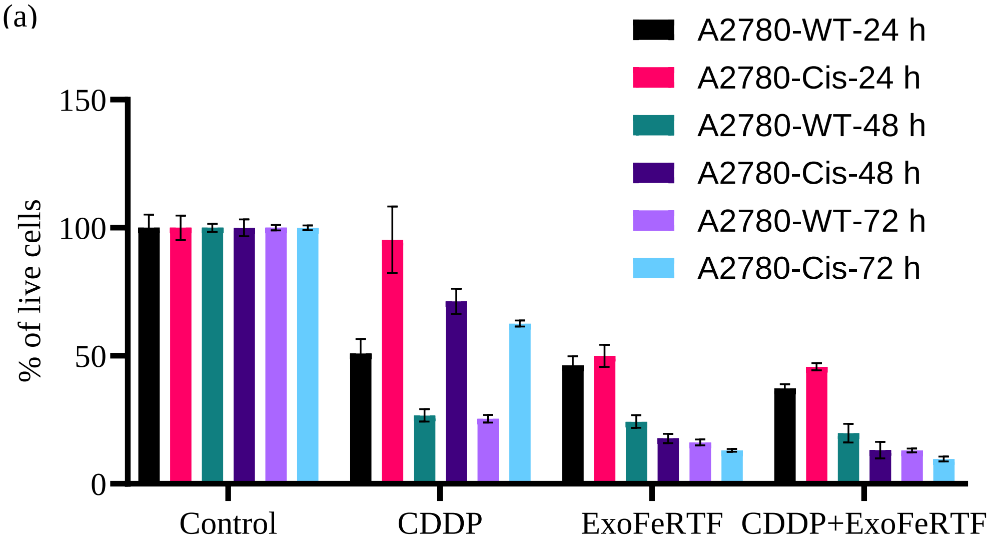


(b)

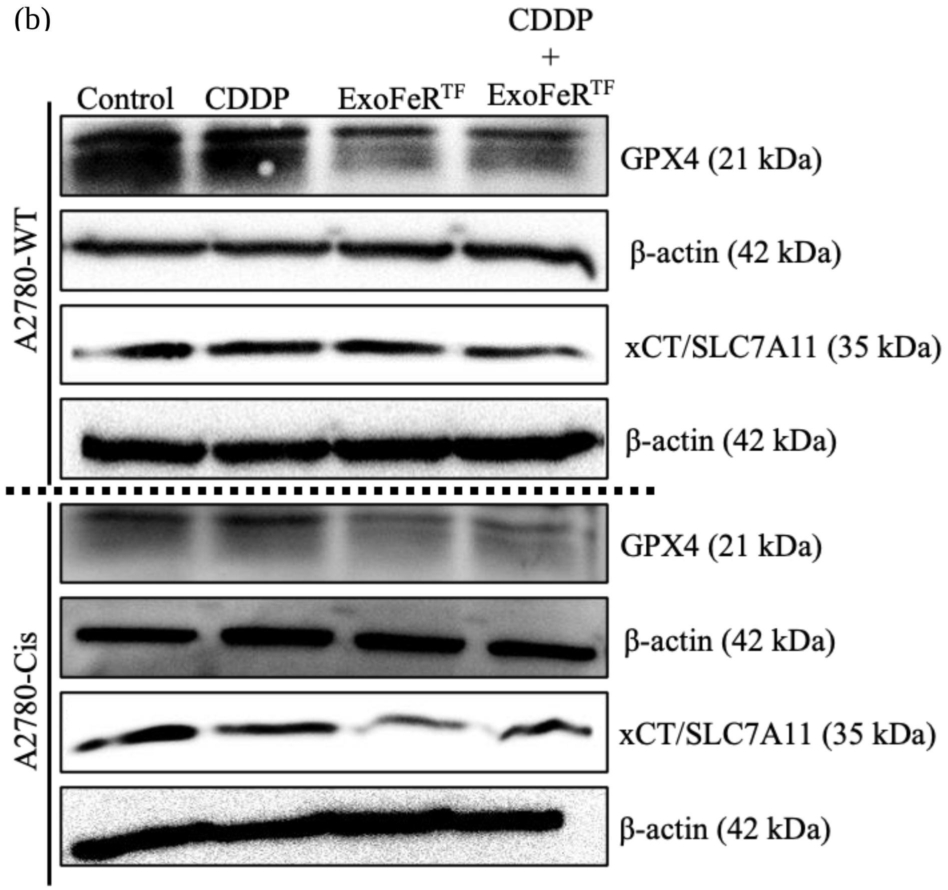


**Supplementary Figure 6. Reversing CDDP resistance with ExoFeR$^{TF}$ Treatment. (a)** Cell viability assays on A2780-WT (parental) and A2780-cis (CDDP resistant) ovarian cancer cells revealed that ExoFeR$^{TF}$, with or without CDDP, induced approximately 70% cell death. However, when combined with CDDP, ExoFeR$^{TF}$ treatment induced about 90% cell death in resistant cells. **(b)** Western blot analysis of ferroptosis markers also demonstrated GPX4 and xCT/SLC7A11 inhibition in the resistant cells following ExoFeR$^{TF}$ and CDDP
